# Structural investigations of the glucan water dikinase 1 mechanism and flexibility

**DOI:** 10.64898/2026.02.06.704335

**Authors:** Thibaud Laffargue, Nina Cooper, David Guieysse, Etienne Séverac, Pascal Mansuelle, Pierre Roblin, Gianluca Cioci, Claire Moulis, Magali Remaud-Simeon

## Abstract

Glucan-water-dikinase 1 (GWD1) plays an essential role in regulating starch metabolism in plants via O-6 phosphorylation of amylopectin. Here, we used biochemical characterization, AlphaFold2 modeling, X-ray crystallography and Small-Angle X-ray Scattering (SAXS) experiments to study its structure and catalytic mechanism. The protein is organized into five domains with two carbohydrate-binding modules (CBMs) at its N-terminal end followed by a central domain, whose structure was solved by X-ray crystallography in open and closed conformations. Next comes the domain carrying the catalytic histidine and the ATP-binding domain. We studied the spatial arrangement of the full enzyme and of several truncated forms by SAXS-driven modeling and identified a pivoting movement of the Histidine domain consistent with the enzyme’s autophosphorylation and subsequent phosphate transfer to a glucan. Our data suggest important residues at the domain interfaces that might assist catalysis and we hypothesize that the second CBM helps maintaining the catalytic domain close to the glucan chain for productive phosphate transfer.

**Graphical abstract:** 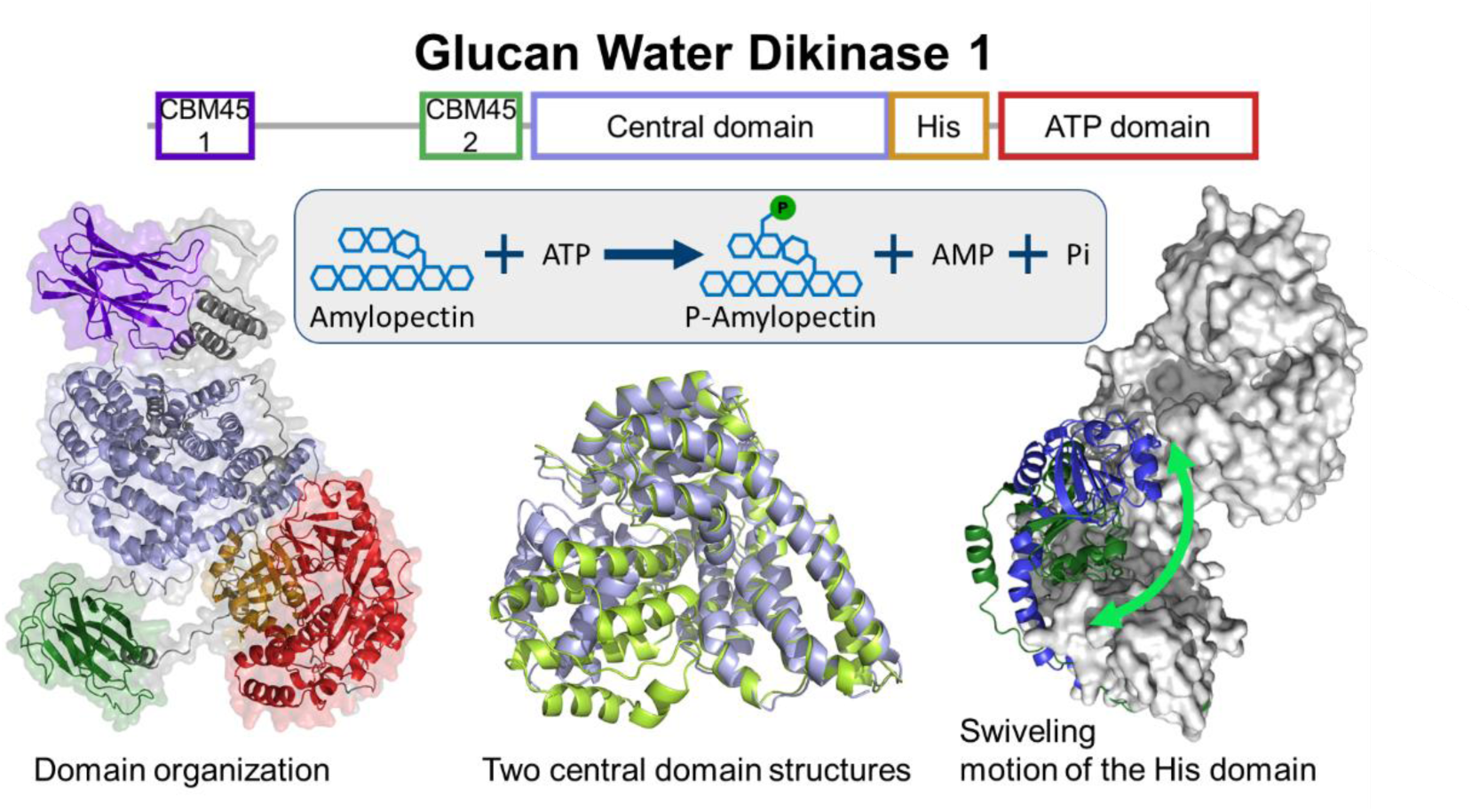

## 1. Introduction

Starch phosphorylation, the only covalent modification occurring naturally to starch, plays a major role in starch metabolism in all higher plants and algae (Blennow, 2015; Compart and Fettke, 2025). Phosphorylation and dephosphorylation are mediated by glucan kinases and glucan phosphatases respectively and the two reactions co-exist in a dynamic process which regulate starch metabolism. Indeed, knock-out of genes encoding glucan kinases or phosphatases affects starch content in addition to phosphorylation levels. (Yu et al., 2001; Kozlov et al., 2007; Kötting et al., 2009; Samodien et al., 2018; Adegbaju et al., 2025). The introduction of phosphate groups in planta mainly occurs in amylopectin and was proposed to destabilize amylopectin double helices and facilitate the access of glucoside hydrolases for degradation (Blennow, 2015; Compart and Fettke, 2025). Starch phosphate content thus varies depending on the plant metabolic state, the producing organisms, and the tissues where starch fulfills different functions (transitory or storage starch). For extensive reviews on the impact of phosphorylation on starch metabolism see (Blennow, 2015; Apriyanto et al., 2022; Compart and Fettke, 2025).

Two different classes of kinases, the glucan water dikinases (GWDs) and the phosphoglucan water dikinases (PWDs) are involved in starch phosphoesterification. GWDs introduce phosphate groups on the free C6-OH of starch whereas PWDs are more specific for C3-OH phosphorylation (Ritte et al., 2002; Baunsgaard et al., 2005; Kötting et al., 2005; Ritte et al., 2006). C6-OH and C3-OH phosphorylation usually accounts for respectively 60-70 % and 30-40 % of total phosphate content in starch (Blennow, 2015). Studies on glucan kinases have been greatly stimulated by the impact of phosphorylation on the physicochemical properties of starch, which reflects on its applications (Compart et al., 2023b; Adegbaju et al., 2025). These enzymes have become prime targets for plant genetic engineering aimed at producing starch with new properties or increasing yields, as illustrated by work on rice, wheat or curcuma starches (Ral et al., 2012; Bowerman et al., 2016; Chen et al., 2017; Wang et al., 2018, 2021). In vitro phosphorylation could also be an alternative for modifying starch or other glucans using these enzymes without resorting to chemicals (Gentry and VanderKooi, 2021; Compart et al., 2023b; Laffargue et al., 2023). All these developments have prompted further research into the catalytic mechanisms of glucan kinases, particularly on GWD1.

GWD1 from *Solanum tuberosum*, StGWD1, is one of the most studied GWD with AtGWD1 from *Arabidopsis thaliana*. It was first characterized by Ritte and collaborators in 2002 (Ritte et al., 2002). Using radioactive ATP labeled with P^33^ either on the β−P^33^ or γ−P^33^ phosphate, it was shown that, StGWD1 catalyzes first its auto-phosphorylation through the transfer of the β-phosphate of ATP to a catalytic histidine (His992 in mature sequence), while releasing the γ-phosphate and AMP. Then, the β-phosphate is transferred to the C6 position of the α-1,4 linked glucosyl units of amylopectin, thus adopting a mechanism characteristic of dikinases (route 1, Figure 1) (Narindrasorasak and Bridger, 1977; Lim et al., 2007; Hsu et al., 2025). Alternatively, StGWD1 can catalyze the transfer of the β-phosphate onto water (route 2, Figure 1). Other studies using P^33^-labeled substrates also reports that multiple phosphotransfer reactions are catalyzed by this enzyme. It was shown that the β-phosphate of ADP and γ-phosphate of ATP can be transferred as a minor reaction onto water, AMP or ADP to yield orthophosphate, ADP or ATP (simplified in Route 3 on Figure 1) (Ritte et al., 2002; Hejazi et al., 2012). Part of the ADP production would thus be the result of AMP phosphorylation, that would occur from ATP or ADP. These reactions would not require the presence of the catalytic histidine, but would involve a second autophosphorylation site, with an amino acid different from the catalytic histidine, able to bind the terminal phosphate of both ADP and ATP and which has not yet been identified (Hejazi et al., 2012). StGWD1 only uses ATP as donor to transfer phosphate to different amylaceous acceptors including amylopectins, glycogens and amyloses of different origins, (Ritte et al., 2002; Mikkelsen et al., 2004; Compart et al., 2023a). The highest phosphate transfer activity was reported on crystalline maltodextrins, a model substrate with a high crystallinity degree considered to be a good mimic of the double helix found in amylopectin (Hejazi et al., 2008, 2009). Recently in vitro assays using different types of starch revealed that StGWD1 can diffuse in the granule and preferentially catalyzes phosphotransfer in the proximity of branching points of amylopectin (Compart et al., 2023a).

**Figure 1:**
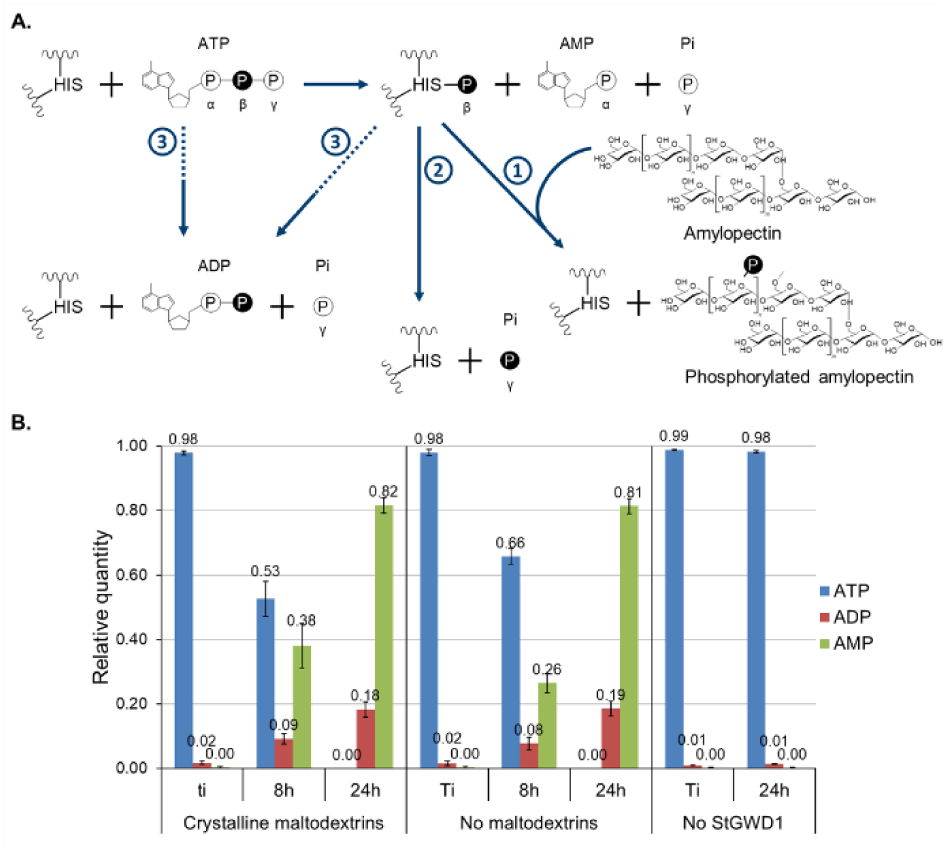
(A) Summary of the various reactions catalyzed by StGWD1. ATP cleavage results in phosphoryl-enzyme intermediate (involving His992), AMP and Pi formation. The β-phosphate linked to His992 is transferred to the α-glucan (route 1) or to water (route 2). The presence of ADP was also reported (Ritte et al., 2002; Hejazi et al., 2012). ADP production would be the result of AMP phosphorylation from ATP or ADP, doesn’t involve the catalytic histidine, but could involve a second autophosphorylation site. (B) Relative quantities of ATP, ADP and AMP over the course of StGWD1 reaction. Reactions on the left were carried out with 1 mM ATP, 1 g.l^-1^ crystalline maltodextrins and 100 µg.ml^-1^ StGWD1 in 50 mM HEPES pH=7.5. Reactions without maltodextrins (middle panel) and controls without enzyme (right panel) were run in parallel. Relative quantities of ATP, ADP and AMP were calculated by dividing the concentration of each compound by the sum of ATP, ADP and AMP concentrations as determined by HPLC analysis, and corresponds to four biological replicates (values at initial and final times), or three biological replicates (values at 8h).

No crystallography structure of GWD1s is available yet. Most of the structure function studies thus rely on biochemical characterization coupled to sequence analyses, mutagenesis studies and generation of truncated forms. StGWD1 partial proteolysis enabled to propose an organization of the protein in five domains (Mikkelsen and Blennow, 2005). The catalytic histidine residue would be located in the 4^th^ domain, and the C-terminal domain was proposed to carry the ATP binding site, due to its sequence homology with the ATP-binding domain of pyruvate, phosphate dikinases (PPDKs) and phosphoenol pyruvate synthases (PEP-synthase, (Lorberth et al., 1998; Yu et al., 2001)). In agreement with these hypotheses, StGWD1 truncated of its 3 first domains retained its ATP cleavage and auto-phosphorylation capabilities (Mikkelsen and Blennow, 2005). These studies suggest that StGWD1 might share a common mechanism with PPDKs and PEP-synthases, in which the histidine domain swivels between the ATP-binding domain and the acceptor-binding domain to transfer the β-phosphate from the ATP to the acceptor molecule. So far, no experimental data came to support the existence of a swiveling motion in StGWD1. Based on sequence alignment, two carbohydrate binding modules from CBM45 family, proposed to form two different domains, were also identified at the enzyme N-terminal extremity (Boraston et al., 2004; Drula et al., 2022). They were suggested to make up the glucan binding domain (Mikkelsen et al., 2006). The function of the remaining domain, located at the center of the protein, is still unknown.

To gain new insights into StGWD1 mechanism, better understand its dynamic and identify molecular determinants important for its action on amylaceous substrates, we initiated structural studies on a recombinant form of StGWD1. We first characterized the activity on crystalline maltodextrins and determined the size of the smallest maltooligosaccharides phosphorylated by StGWD1. Since the full-length enzyme was refractory to crystallization, we generated truncated forms aided by AlphaFold2 prediction, we characterized their activities and structural features using X-ray crystallography and small angle scattering (SAXS). This led to the resolution of the 3D-structure of the central domain of StGWD1. We investigated the spatial arrangement and organization of the different modules of the protein and acquired information on the protein dynamics. Our findings are consistent with a movement of the histidine domain that is required for productive catalysis, as already observed for other dikinases, and also suggest a different role for each of the CBM45 modules in catalysis. Our work sheds new light on the complex mechanism of starch phosphorylation mediated by GWD1.

## 2. Results

### 2.1 Phosphorylation of crystalline maltodextrins with StGWD1

StGWD1 (155 kDa) was recombinantly expressed in *E. coli* without its transit peptide and the purified protein obtained after anion-exchange and size-exclusion chromatography with a yield of 125 mg/L_culture_ was used throughout the rest of our study (Figure S1). We tested the activity of StGWD1 in the presence of ATP and crystalline maltodextrins having a degree of polymerization (DP) ranging from 3 to 50 (Figure 1), known to be efficient acceptor (Hejazi et al., 2008). Control reactions without glucan or enzyme were also carried out. In the enzyme-free control, ATP was not significantly degraded and remained stable under the test conditions. The ATP consumption was followed under the course of the reaction and found to be linear during the first 8 hours (Figure S2). After 8 hours, the relative quantity of AMP was higher in the presence of maltodextrins (38 % vs 26 %), corresponding to higher ATP depletion (53 % vs 66 %). ATP consumption with and without maltodextrins, estimated after 8 hours reaction, was 10 and 7 nmol.min^-1^.mg^-1^, showing StGWD1 is activated in the presence of the α-glucan, as described in other studies (Ritte et al., 2002; Hejazi et al., 2012). After 24h, ATP was totally consumed and converted to ADP and AMP, representing 18 % and 82 % of the initial ATP concentration, respectively.

Capillary IC-MS revealed the presence of mono-phosphorylated compounds of DP5 to DP35, di-phosphorylated products of DP12 to 36 and tri-phosphorylated products of DP19 to 30 (Figure 2A, and S3 to S6). Products of higher DPs might be present, but not detectable with our system. Using classic IC-MS, we could detect shorter mono-phosphorylated malto-oligosaccharides (DP3 to DP4). To determine total phosphate incorporation, we submitted StGWD1-modified maltodextrins to total acid hydrolysis, and quantified the glucose and glucose-6-phosphate (G6P) using HPAEC-PAD and IC-mass spectrometry, respectively. After 24 hours of reaction, 20.2 µM of G6P were detected, corresponding to a degree of substitution (DS) of 0.0033.

**Figure 2:**
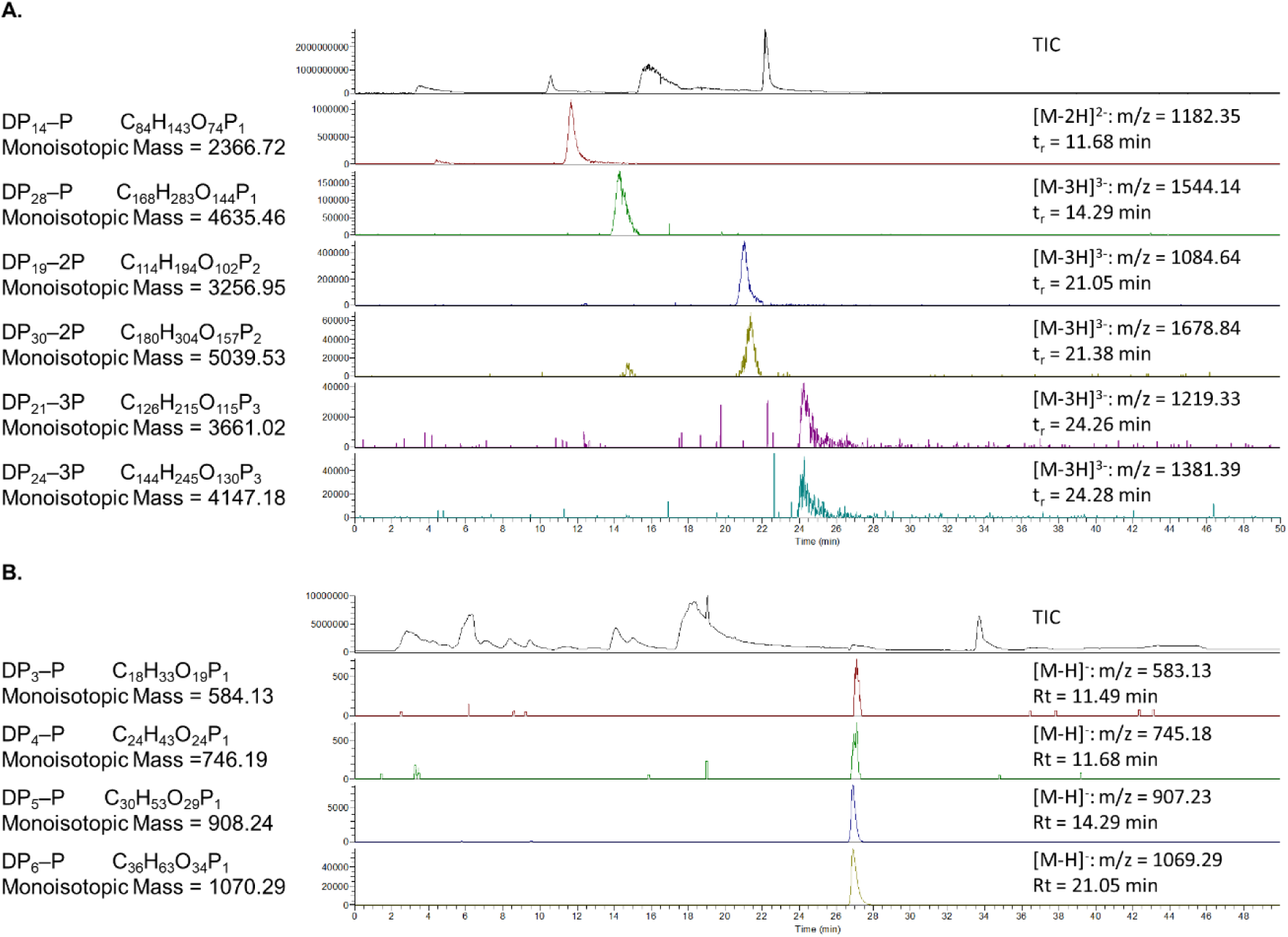
Phosphorylated maltodextrins detected in the reaction media of StGWD1 with crystalline maltodextrins and ATP. (A) Extracted ion chromatograms (EIC) of 7 mono-, di- or tri-phosphorylated products (selected among all the ones detected) detected by capillary IC-MS (see figures S3 to S6 for more details). (B) Phosphorylated maltooligosaccharides of DP3 to DP6 detected by IC-MS. Monoisotopic masses used are indicated on the left, and extracted masses as well as retention time (Rt) on the right. Total Ion Count (TIC) is presented at the top.

### 2.2 StGWD1 a large monomer stabilized by AMP-PNP

SEC-MALS experiments showed that StGWD1 is a monomer in solution with an estimated molecular weight of 147 kDa, close to the theoretical value (154.8 kDa, Figure S1), and a hydrodynamic diameter of 6.0 nm measured by Dynamic Light Scattering results (DLS). The thermal denaturation profile determined without ligand by nanoDSF shows two inflection points, corresponding to two melting temperatures (Tm_1_ of 54.6 °C and Tm_2_ of 61.2 °C) and suggesting that different domains could unfold at different temperatures (Figure S7). The addition of carbohydrate ligands as well as ATP, ADP, and AMP in the presence of MgCl_2_ did not affect the denaturation profile (Table S1). Adenylyl-imidodiphosphate (AMP-PNP), a non-hydrolysable ATP analogue, was found to be the best stabilizing ligand. It was used to setup crystallization assays for the full-length StGWD1, which however failed to give any starting condition probably due to inherent enzyme flexibility. Therefore, we turned to molecular modelling and we generated structure predictions using AlphaFold2 (AF2).

### 2.3 Description of StGWD1 model

The AF2 model shows 5 domains (Figure 3 and Figure S8 for more details) predicted with high confidence score (pLDDT > 70), connected by flexible linkers (pLDDT < 70). The first two domains, CBM45_1 (aa 11 to 129) and CBM45_2 (aa 343 to 464) belongs to CBM45 family according to CAZy classification (Drula et al., 2022), as previously reported by Mikkelsen and collaborators (Mikkelsen et al., 2006). They are formed by 11 and 9 β-strands for CBM45_1 and CBM45_2, respectively (Figure S8) and contain the conserved Trp residues shown to be important for starch binding (Mikkelsen et al., 2006; Adegbaju et al., 2020).

**Figure 3:**
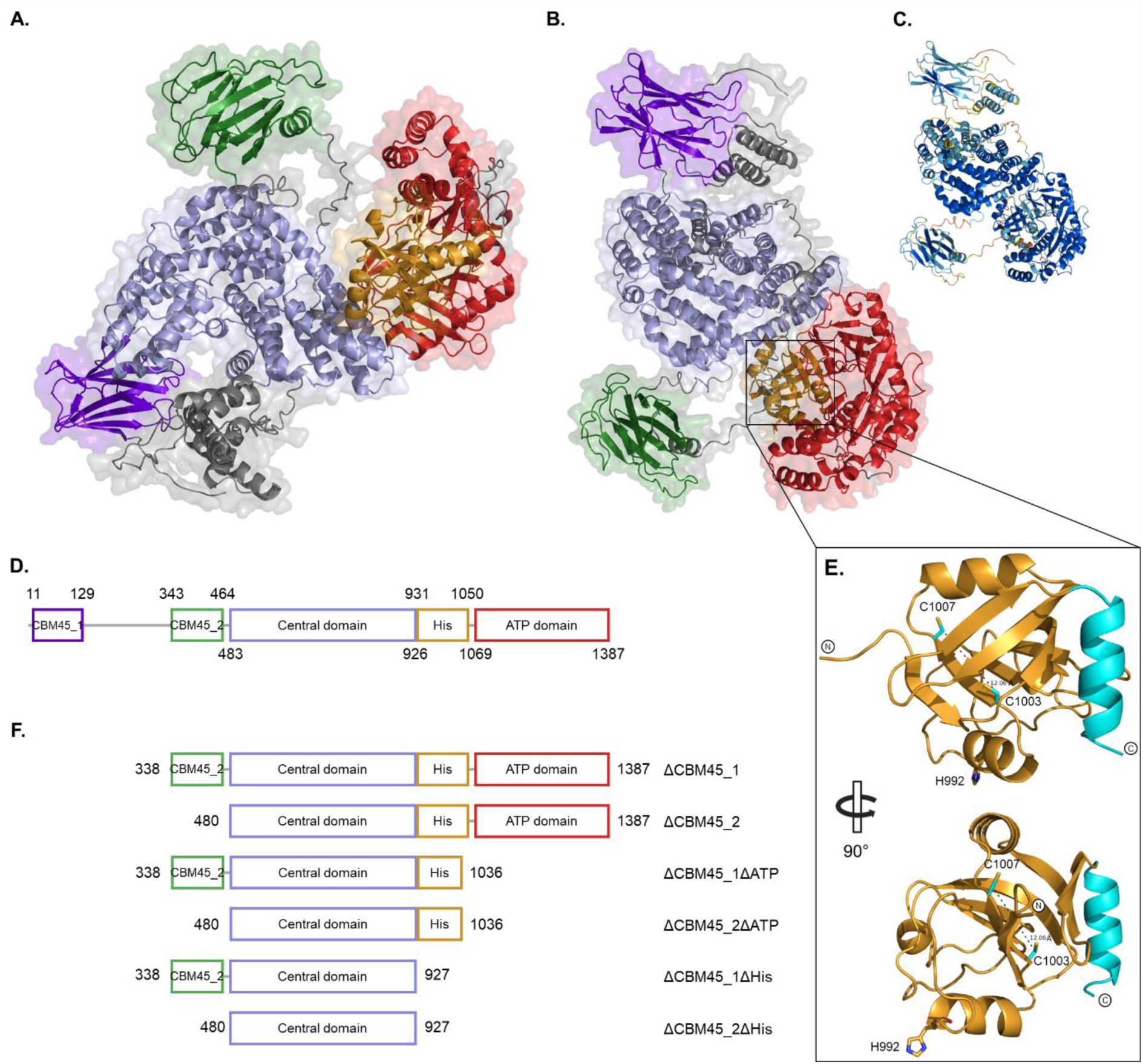
(A) and (B) Two views of the AlphaFold2 model of StGWD1, with each domain colored as presented on panel D. (C) coloration following AlphaFold2 confidence score (pLDDT). Orange: pLDDT < 50; yellow: pLDDT > 50; light blue: pLDDT > 70; dark blue: pLDDT > 90. (D) Schematic representation of StGWD1. Squares represent folded domains; gray lines represent unfolded regions. (E) Model of the histidine domain of StGWD1, with the catalytic H992, C1003 and C1007 shown as sticks. The α-helix that could be part of the linker is shown in light blue (F) Truncated forms of StGWD1 designed based on the domains provided by the AlphaFold2 model.

The third domain or “central domain” (aa 483 to 926) was crystallized and will be described in detail later. The fourth domain (aa 931 to 1050), called the histidine domain (his domain), consists of 3 α-helices and 7 β-strands and contains the solvent-exposed catalytic histidine (His992, Figure 3E). An additional α-helix (low pLDDT) at the C-terminus is further away and probably part of the linker between the histidine domain and the ATP-binding domain (aa 1069 to 1387). The latter folds into 10 α-helices and 12 β-strands and is homologue to the ATP-binding domain of *Flaveria trinervia* PPDK (FtPPDK), as initially suggested from sequence analysis (Lorberth et al., 1998; Yu et al., 2001), and to *Listeria monocytogenese* Rifampin Phosphotransferase (LmRPH), an enzyme responsible for the phosphorylation of the antibiotic rifampicin (Qi et al., 2016; Stogios et al., 2016).

In the different models proposed by AF2, the position of CBM45_1 varied considerably relative to the rest of the structure which forms a more compact object. Notably, the His and ATP-binding domains closely interact allowing the catalytic histidine to insert in the putative ATP-binding site. Not surprisingly, the 5 domains of the StGWD1 model agreed with the fragments that we obtained by partial proteolysis of the enzyme (Figure S9).

### 2.4 The CBMs influence ATP consumption

From the AF2 model, we designed, expressed and purified six truncated forms (Figure 3F). For ΔCBM45_1 and ΔCBM45_2, the two forms active on ATP, the relative quantities of ATP, ADP and AMP formed after 8 h of incubation are quite similar (Figure 4). Removing the first CBM45 leads to a decrease in ATP consumption and also eliminates the maltodextrin-dependent increase in activity that is observed for full-length StGWD1. At 24 h, ΔCBM45_1 and ΔCBM45_2 converted 98 % and 75 % of ATP, respectively. This suggest that ΔCBM45_2 is less stable than ΔCBM45_1.

**Figure 4:**
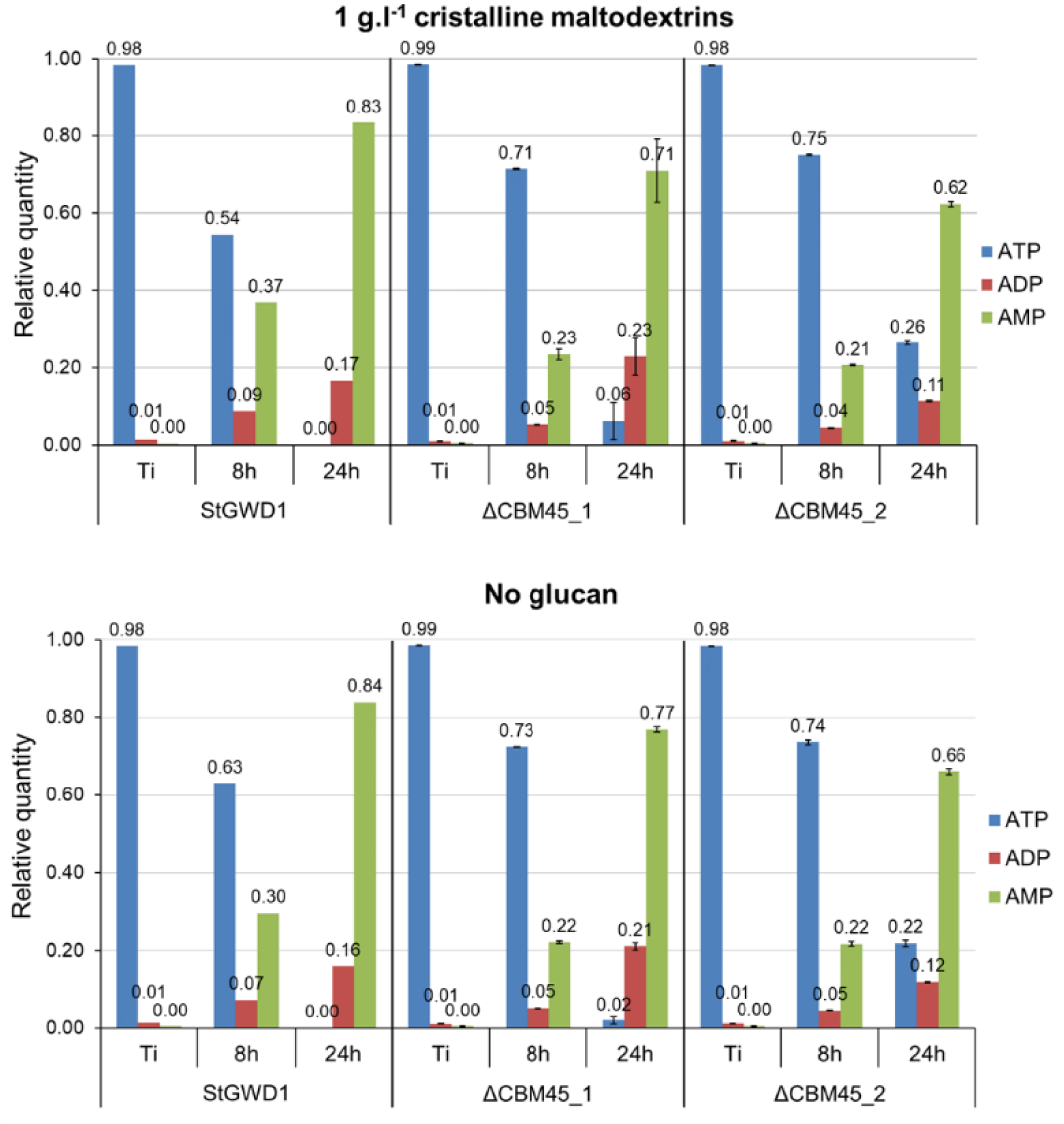
Relative quantities of ATP, ADP and AMP over the course of StGWD1, ΔCBM45_1 and ΔCBM45_2. Reactions were carried out with 1 mM ATP and 670 nM enzyme in 25 mM HEPES pH=7.5 in presence (top panel) or absence (bottom panel) of 1 g.l^-1^ crystalline maltodextrins. Relative quantities of ATP, ADP and AMP were calculated by dividing the concentration of each compound by the sum of ATP, ADP and AMP concentrations as determined by HPLC analysis, and corresponds to three technical replicates (for ΔCBM45_1 and ΔCBM45_2).

### 2.5 Crystal structure of the central domain of StGWD1

All the truncated forms were submitted to crystallization trials with or without appropriate ligands (maltooligosaccharides and/or AMP-PNP/Mg^2+^). We obtained crystals only for ΔCBM45_2ΔHis, called the central domain, and its 3D structure was solved by molecular replacement at 3.0 Å resolution (Figure 5 and Table S2). This domain is composed of 20 α-helices arranged in three clusters, and adopts a reverse “U” shape: cluster 1 (helices 1 to 5) at the top of the reverse “U”, and cluster 2 (helices 6 to 11) and cluster 3 (helices 12 to 20) on either side. Interestingly, we obtained two conformations, termed “open” and closed”, both adopting an overall structural arrangement with no clear equivalent exists in the PDB. In the open form (Figure 5A), which is highly similar to the AF2 prediction, clusters 1 and 3 interact strongly with each other, through contacts between helices 1, 3, 5 and helices 15, 18 and 19, respectively. Cluster 2 is more distant and has very little contacts with clusters 1 and 3. The two helices 6 and 11 of cluster 2 are side by side, creating a single “attachment point” that connects it to cluster 1 via the hinge loop between helix 6 and helix 5, and to cluster 3 via the loop between helix 11 and helix 12. The position of cluster 2 is thus such that a central groove is formed. The electrostatic potential map showed some patches of negative charges on the surface of this domain but their biological significance is unclear (Figure S10). The second crystal form was obtained in a different crystallization condition and diffracted at lower resolution of 3.25Å (Figure 5C and Table S2). In this structure, the central domain has a “closed” conformation that is likely due to the flexibility of the hinge loops linking clusters 1-2 and 2-3, causing cluster 2 to tilt and come into tight contact with cluster 3, completely closing the central grove present in the open form (Figure 5D). Interestingly, small angle X-ray scattering (SAXS) is not able to clearly distinguish one conformation in solution, suggesting that both may be biologically relevant (Figure S11).

**Figure 5:**
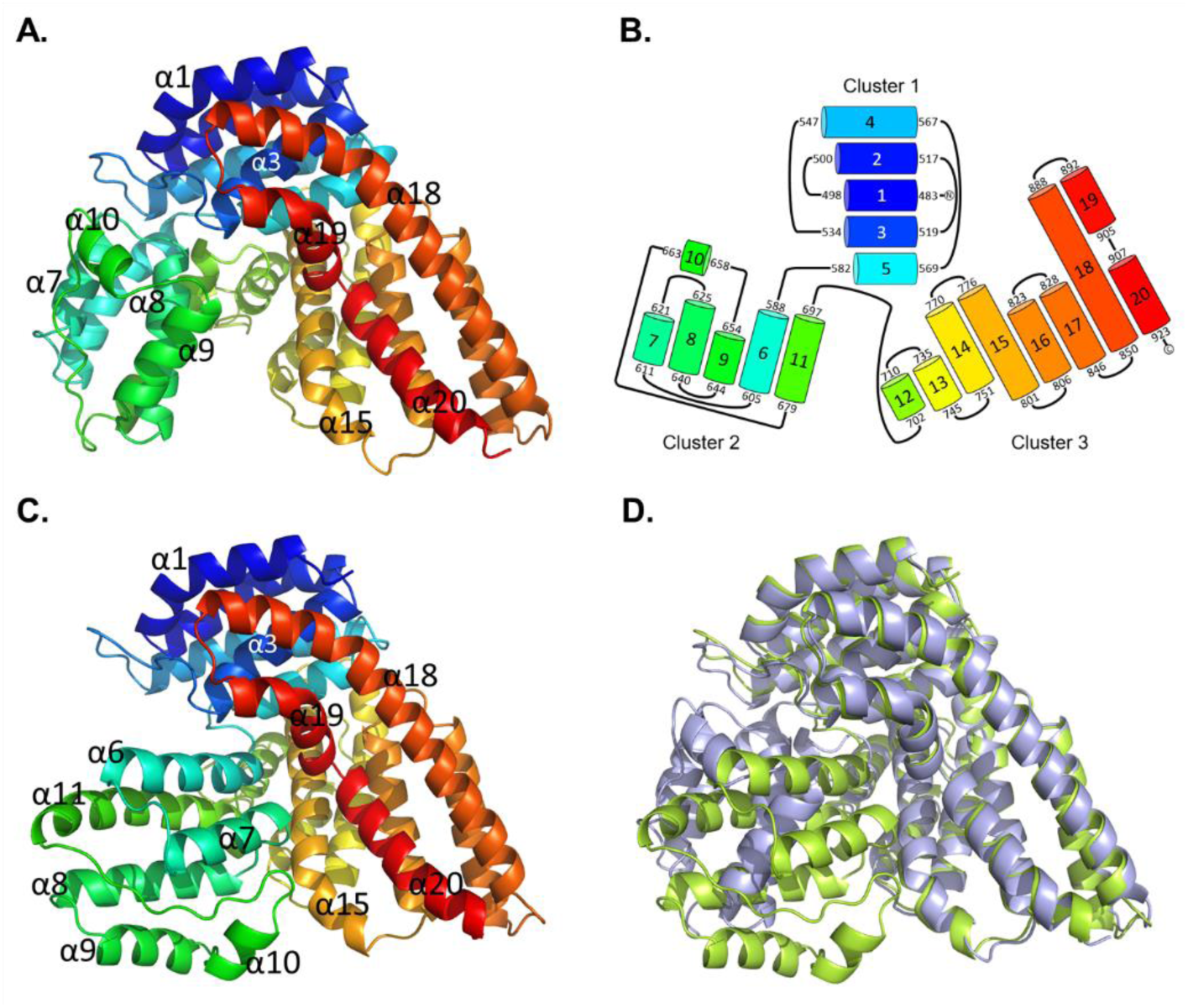
Crystal structures obtained for the central domain of StGWD1 (A) Open conformation of the central domain (PDBID 9HA9), where cluster 2 does not interact with cluster 3, creating a central groove between them. (B) Topology diagram. The three clusters of α-helices are shown, with their relative position and orientation in space, colored as in A. (C) Closed conformation of the central domain (PDBID 9QVI), where cluster 2 is in contact with cluster 3, closing the central groove. (D) Superposition of the open (mauve) and closed (green) structures obtained for the central domain, highlighting the tilting of cluster 2.

### 2.6 Substrate induced movement of the histidine domain

The full-length StGWD1 and all the other truncated forms were analysed by SAXS. The Rg and the Dmax (Table S3, Figure S12) are positively correlated to the molecular mass of the objects studied. On the Kratky plot obtained for the full-length protein the major peak has a sRg > √3 and a shoulder can be seen at sRg ∼ 6. The pair distribution function shows a large asymmetric peak and a shoulder slowly going to zero at higher distances with a Dmax of the particle of ∼200 Å (Figure S12). All these elements are characteristic of multidomain proteins with flexible linkers (Rambo and Tainer, 2011). The DADIMODO program (Rudenko et al., 2019) was used to perform rigid-body fitting of the full-length AF2 model against the SAXS data. Flexibility was allowed for the linker regions between all the structured domains but the relative position of the ATP-binding domain to the central domain was kept fixed. The resulting model fitted the SAXS data well, with a χ² of 2.77 (Figure 6A). In this model, the CBM45_2 is sitting in the central globular part, half-way between the central and the ATP-binding domain and close to the position proposed by AF2. In contrast, the CBM45_1, along with the unstructured part connecting the two CBMs, was moved further away in agreement with the elongated character of the protein. The DADIMODO model also superposes reasonably well the *ab-initio* envelop generated from the SAXS data (Figure 6A). The envelop presents a big globular domain, with a thinner long “arm” that could well represent the average position of the CBM45_1 along with the partially unfolded domain connecting it to the rest of the protein.

**Figure 6:**
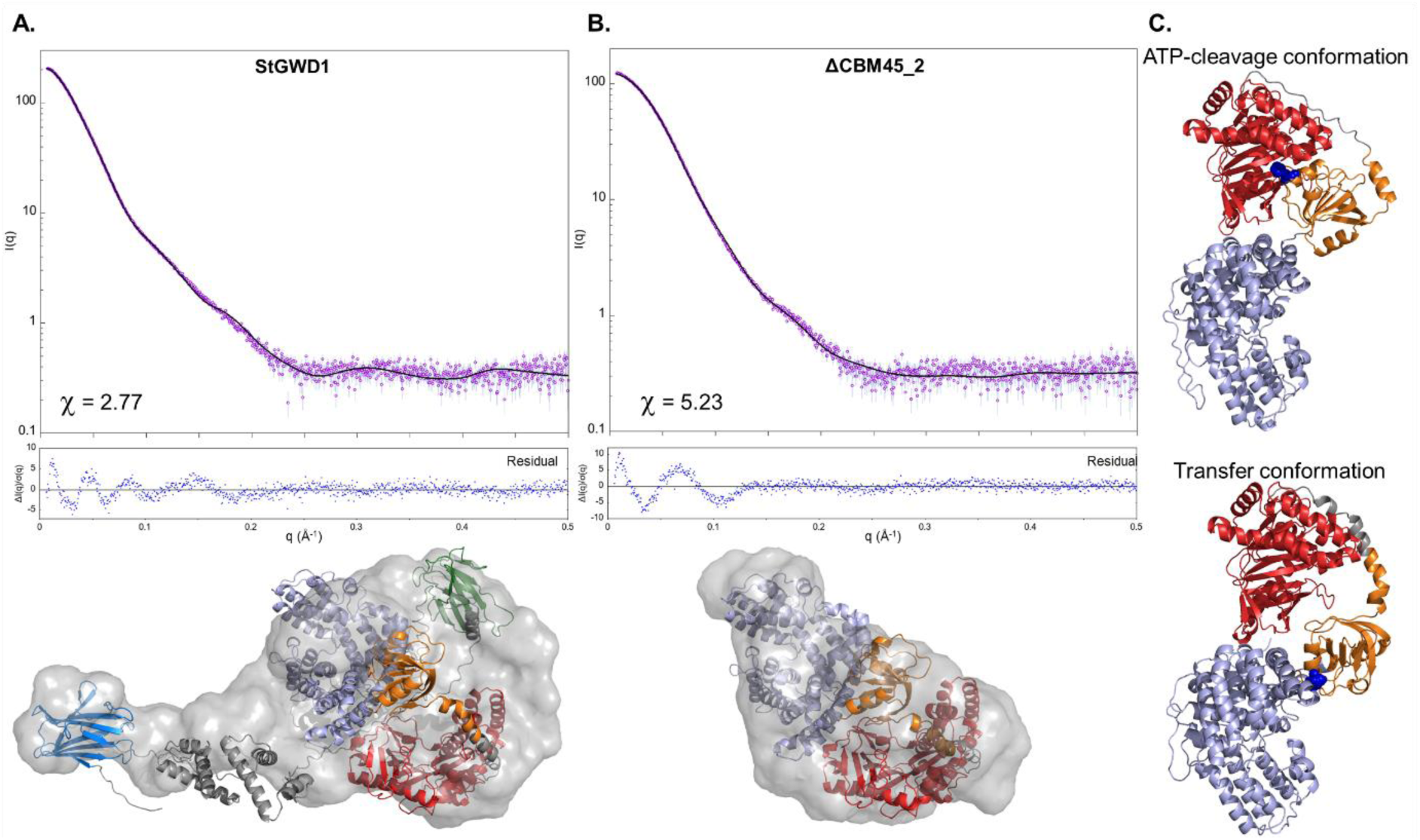
Best fitting model of (A) StGWD1 and (B) dCBM2 obtained using DADIMODO, superposed with one representative ab-initio envelop (bottom). Theoretical data calculated for the models (light blue curve) are compared with the experimental data (blue points with error bars). (C) Two conformations of ΔCBM45_2. Top: ATP-cleavage conformation, with the central, histidine and ATP-binding domains as seen in the AF2 model of full-length StGWD1. Bottom: transfer conformation, with the histidine domain flanking the central domain. The catalytic histidine is shown in blue spheres. Purple: CBM45_1; Green: CBM45_2; Mauve: Central domain; Orange: Histidine domain. Red: ATP-binding domain. Gray: linker regions.

Regarding the truncated forms, all the Kratky plots, except for ΔCBM45_1ΔHis for which we could not obtain exploitable data, revealed a main peak with an sRg almost equal to 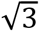 indicating that these proteins are more globular objects than the full-length StGWD1 (Figure S12). A change is clearly visible between the full-length protein and ΔCBM45_1, suggesting that the N-terminal region contains most of the unfolded and flexible regions of the protein. Moreover, the proteins truncated of their N-terminal parts, *i.e.* ΔCBM45_1 and ΔCBM45_2, have very similar Rg and Dmax, with Dmax being ∼30 % lower than the Dmax of StGWD1 (Table S3). This confirms that the CBM45_2 should sit fairly close to the globular part of the protein whereas the CBM45_1 should be located further.

To attempt rigid body fit of the truncated forms, we first generated AF2 models for each of them. Surprisingly, for ΔCBM45_2ΔATP, AF2 predicted a new conformation in which the histidine domain is flanking the central domain and the catalytic histidine pointing towards the central groove. Starting from this, we generated new models of the full-length protein and its truncated variants in two different conformations: (i) the “ATP-cleavage” conformation, directly derived from the AF2 model of the full-length protein, in which the histidine domain is close to the ATP-binding domain to attack ATP and (ii) the “transfer” conformation, in which the histidine domain has moved towards the central domain, with the catalytic histidine pointing to the central domain in a position likely to be more favorable for transfer to the acceptor (Figure 6C). Rigid body fit on the full-length enzyme, ΔCBM45_1, ΔCBM45_2 and ΔCBM45_2ΔATP was performed using DADIMODO as described above, starting from either the “ATP-cleavage” or the “transfer” conformation. All the models converged to a position of the histidine domain that is close to the transfer conformation (Figure 6A and 6B, S11 and S13), regardless of the starting conformation used, suggesting that in solution the His domain has a resting position near the central domain, and moves towards the ATP-binding domain upon nucleotide binding. To test this hypothesis, SAXS data of ΔCBM45_2 were collected in presence and absence of AMP-PNP/Mg^2+^, whose stabilizing effect on the construct was verified (Figure S14). The OLIGOMER software (Konarev et al., 2003) calculated that in absence of AMP-PNP/Mg^2+^ the ATP-cleavage conformation contributed to 21 % of the observed total scattering, while the transfer conformation accounted for 79 % (Figure S15). In the presence of AMP-PNP/Mg^2+^, a clear switch occurs, with the ATP-cleavage conformation reaching 100 % and the transfer conformation dropping to 0 %. These data support that a clear conformational shift of the histidine domain occurs from the central domain to the ATP-binding domain in presence of ATP, as demonstrated for other dikinases (Lim et al., 2007; Stogios et al., 2016).

## 3. Discussion

To get insight in the mechanism of StGWD1 and as a premise to engineering work with the aim of improving its efficiency for the phosphorylation of α-1,4 glucans and extending its acceptor substrate specificity to different α-glucans, we produced StGWD1 recombinantly, characterized its action on crystalline maltodextrins and initiated structure-function studies. The results obtained from X-ray crystallography and SAXS experiments, coupled with AF2 modelling, allow us to propose a solution model for the overall structure and for its truncated forms, and to identify important features for its biological activity.

### 3.1 Crystalline maltodextrin phosphorylation approaches in vivo starch phosphorylation levels

Phosphorylation reactions were performed with 1 mM ATP, 1 g/L crystalline maltodextrins, and 0.1 g/L enzyme, allowing direct quantification of ATP consumption, AMP and ADP formation, and determination of the degree of glucan phosphorylation without the use of radioactive substrates. The ATP concentration was selected to maximize glucan phosphorylation and enzyme loading to ensure complete ATP consumption within 24 h. Effectively, all ATP was consumed after 24 h, and only 20 µM phosphate (representing ∼2% of the total ATP hydrolyzed) was incorporated into maltodextrins. This indicates that ATP cleavage to AMP without concomitant glucan phosphorylation, likely occurring primarily via enzyme self-phosphorylation (route 2), appears to be the predominant reaction. Therefore, the initial ATP pool is unlikely to limit glucan phosphorylation. The degree of maltodextrin phosphorylation obtained was of 0.0033 corresponding to 1 phosphate per 300 glucosyl units, which is close to the level observed *in vivo* in potatoes (0.005 or 1 phosphate per 200 glucosyl units (Blennow et al., 1998)).

ATP consumption rates over 8 h were 10.0 and 7.0 nmol·min⁻¹·mg⁻¹ StGWD1 in the presence and absence of crystalline maltodextrins and using 1mM ATP, respectively. These values are within the range reported previously: 50 and 10 nmol·min⁻¹·mg⁻¹ with or without crystalline maltodextrins, respectively, at 10 µM ATP (Hejazi et al., 2012). Notably, StGWD1 activity is less increased in the presence of crystalline maltodextrins. This may reflect ATP-dependent inhibition particularly in the presence of glucan as ATP concentration was higher in our study. Consistent with this, increasing the ATP concentration to 3 mM reduced ATP consumption rate over 8 h to 3.8 nmol·min⁻¹·mg⁻¹ StGWD1 (Figure S2).

Without glucan, we obtained an ADP/AMP ratio of 0.23 which is lower than those previously reported by Hejazi et al., 2012 (0.8 to 2 depending on reaction advancement, and using radioactive ADP/radioactive Pi, which reflects the ADP/AMP ratio). The rate of the different phosphate transfer reaction thus strongly depends on the reaction conditions. To allow proper comparison and a better understanding of the catalytic mechanism, the concentrations of all products formed over time, including ATP, ADP, AMP as well as Pi (which was not determined in our study), should be systematically quantified in addition to the phosphate incorporated into the glucan.

### 3.2 StGWD1 is a multidomain protein and its intrinsic flexibility supports catalysis

SEC-MALS analyses of StGWD1 confirmed that the protein is monomeric rather than dimeric, as reported in the review from Compart at al., 2025. The AlphaFold2 model predicted a five-domain organization consistent with that also reported in Compart at al., 2025, but different from the initially proposed organization, that was deduced only from protein partial proteolysis (Mikkelsen and Blennow, 2005). Our SAXS analysis goes further in the structural characterization, showing that the N-terminal part is responsible for the non-globularity of the protein. CBM45_1 is indeed connected to the protein through a long linker and SAXS-driven modelling positioned it further away from the catalytic core. These elements strongly suggest that the position of the first CBM45 is therefore not fixed, but that it can move around the enzyme. We can hypothesize that this domain would help the enzyme locate the starch granule and guide it to accessible sites. In contrast, the second CBM45, being less mobile and located closer to the central domain, would ensure the proximity and the spatial orientation needed to catalyze glucan phosphorylation.

It is proposed that StGWD1 preferentially phosphorylates close to the branching points of amylopectin and therefore must penetrate the surface of the starch granule surface to access its substrate (Compart et al., 2023a). Given the dense packing of double helices at the granule surface, accommodation of such a large enzyme between them is likely to be sterically challenging. Although the intrinsic flexibility of StGWD1 certainly helps enzyme penetration anywhere in the granule surface, it cannot be excluded that only less densely packed regions are accessible to the enzyme. This would imply that accessible phosphorylation sites are rare, as previously suggested (Compart and Fettke, 2025), in particular if the enzyme needs to reach glucan regions close to branching points. In addition, the presence of soluble glucan chains, whether bound to the granule surface or free in solution was shown to reduce StGWD1 phosphorylation activity (Mahlow et al., 2014). We suggest that soluble chains act as competitive binders to the enzyme CBMs, thereby hindering the enzyme progression into the granule surface, and explaining the reduced activity.

Regarding the central domain, two distinct crystal structures were obtained, in which cluster 2 is either positioned close to cluster 3 or located further away, thereby forming a central groove. SAXS data indicate that both conformations might be equally possible in solution, making it difficult to comment on the biological relevance of this conformational change. Nevertheless, the latter may be useful to penetrate the starch granule surface and moreover to precisely accommodate the glucan substrate, further supporting the notion that StGWD1 is an intrinsically flexible enzyme.

### 3.3 StGWD1 shares distant structural homology with other dikinases

Although the primary-sequence arrangement of the domains (ATP-binding, His and central domain or its equivalent) differs dramatically among StGWD1, PPDKs, and LmRPH, their His and ATP-binding domains exhibit interesting common conserved structural features.

The ATP-binding domains of StDWD1 and PPDKs are homologous and differ primarily due to the presence of an additional subdomain in PPDKs. Interestingly, StGWD1 ATP-binding domain aligns much better with that of LmRPH (Figure 7A). Most of the residues interacting with AMP-PNP/Mg^2+^ in the complexes LmRPH:AMP-PNP (PDBID 5HV3, (Qi et al., 2016)) and FtPPDK:AMPPNP (PDBID 5JVL, (Minges et al., 2017)) are conserved and superimpose well with their counterparts in our model. Although residues E1225, I1226, and I1227 are not strictly conserved among the three enzymes, their backbones overlap allowing conserved hydrogen-bonding interactions with the purine ring (Figure 7B). Overall, the conserved ATP-binding site architecture suggests a shared mechanism for ATP binding.

**Figure 7:**
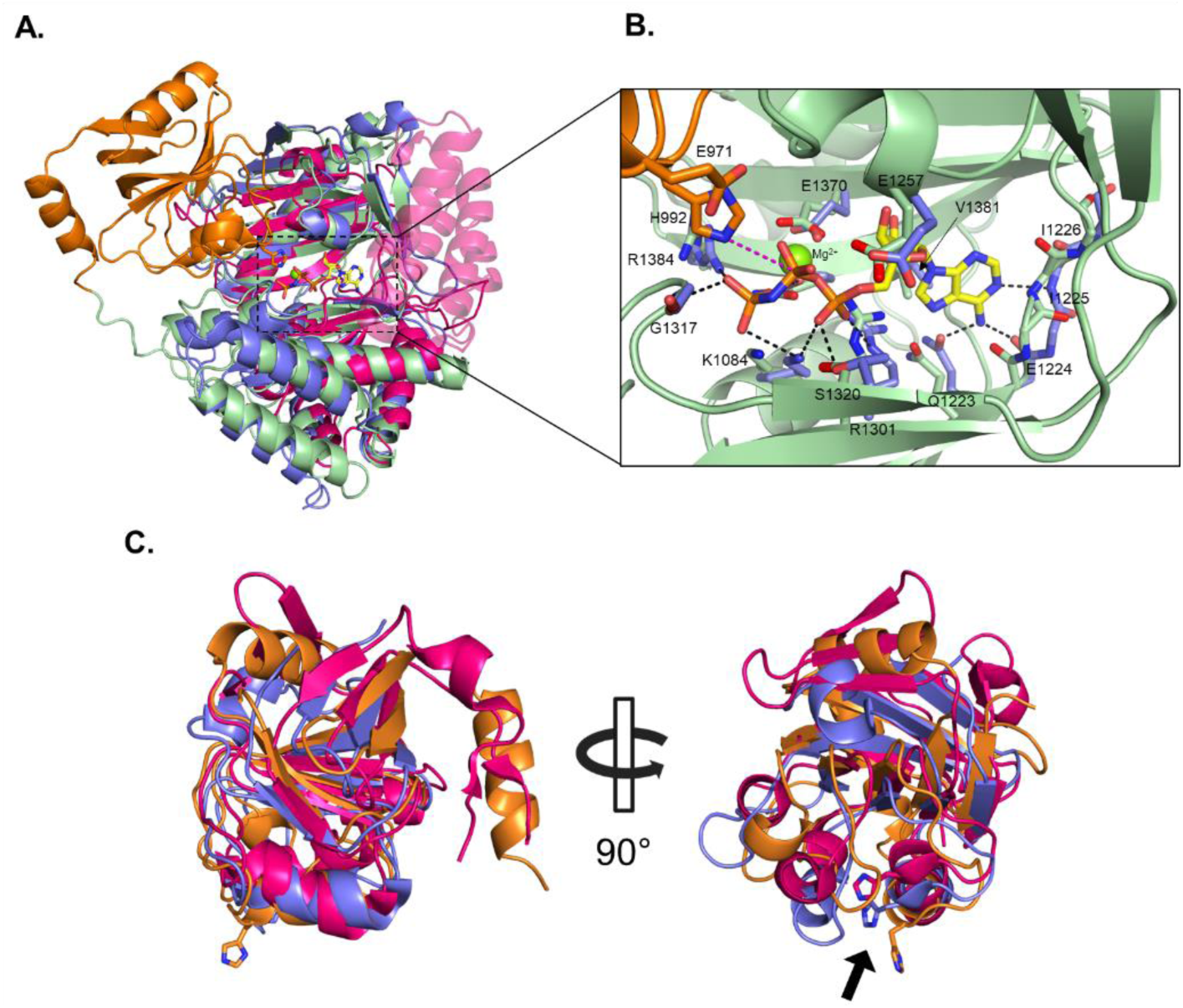
Structural basis for StGWD1 autophosphorylation. (A) Structural comparison of ATP-binding domains of StGWD1 (AF2 model, green), FtPPDK (PDBID 5JVL, pink), and LmRPH (PDBID 5HV3, blue). The AMP-PNP/Mg^2+^ bound in LmRPH structure is shown as sticks. The His domain of StGWD1 is also shown (orange) along with the catalytic histidine H992. (B) Zoom on the ATP-binding domain of StGWD1 (AF2 model, green), superposed with the AMP-PNP/Mg^2+^ bound in the LmRPH (PDBID 5HV3), along with its interacting residues (blue). The His domain of StGWD1 is shown in orange with the catalytic histidine H992 and E971 shown as sticks. Labeled residues refer to StGWD1. (C) Structural comparison of His domains of StGWD1 (our AF2 model, orange), FtPPDK (PDBID 5LU4, pink) and LmRPH (PDBID 5HV3, blue). The position of catalytic histidines (H992 in StGWD1) are shown as sticks and indicated with an arrow.

The His domain of StGWD1 distantly resembles those of LmRPH, or FtPPDK (Figure 7C), the similarity being limited to the common conserved fold, which exposes the catalytic His on the same short helix (StGWD1 numbering: H992-G1000). In our AF2 model, this helix interacts closely with the ATP-binding domain, positioning the catalytic histidine H992 close to the β-phosphate, providing the structural basis for autophosphorylation (Figure 7B). Based on this model, phosphorylation should occur on the Nε atom of the histidine and could be assisted by E971, present in the His domain, which would likely polarize the histidine, to favor nucleophilic attack on the phosphate. Additionally, AMP-PNP binding to LmPRH induces the formation of an α-helix and rigidification of a previously flexible loop, both conserved in StGWD1 (aa 1256–1260 and aa 1306–1315, respectively). This was proposed to create steric repulsion between the histidine and the ATP-binding domain promoting the swiveling motion (Qi et al., 2016). A similar mechanism can be suggested for StGWD1.

### 3.4 Evidences of swiveling motion of StGWD1 His domain conserved through evolution

Our structural data supports that in the absence of ATP, the His domain of StGWD1 adopts a resting state close to the central domain, and that it moves towards the ATP-binding domain to perform autophosphorylation, as described for other dikinases (Stogios et al., 2016; Minges et al., 2017). Upon autophosphorylation, the His domain moves back to the central domain to recreate the proper interface for glucan binding and phosphate transfer. Coevolution may have occurred at this interface, prompting us to search for conserved residues among StGWD1 homologues (Figure 8). The interface involves residues from helices 18, 19 and 20 of the central domain, and residues from the first β-strand, the short helix containing the catalytic His, and a loop (aa 986 to 991) of the His domain. The conservation analysis mainly highlights a highly conserved hydrophobic pocket formed between helices 18 and 20 (central domain), that interacts with highly conserved V989 and L990 of the His domain. Additionally, several conserved polar residues have the potential to form hydrogen bonds (e.g. R923 with W931 backbone, Figure 8A and 8C). Overall, this analysis further supports the phosphate transfer conformation of the His domain and the proposed swiveling motion. The movement of the His domain needs that the ATP-binding and central domains remain in a fixed position relative to each other. This could be ensured by the stable interface between the two domains described in Figure S16.

**Figure 8:**
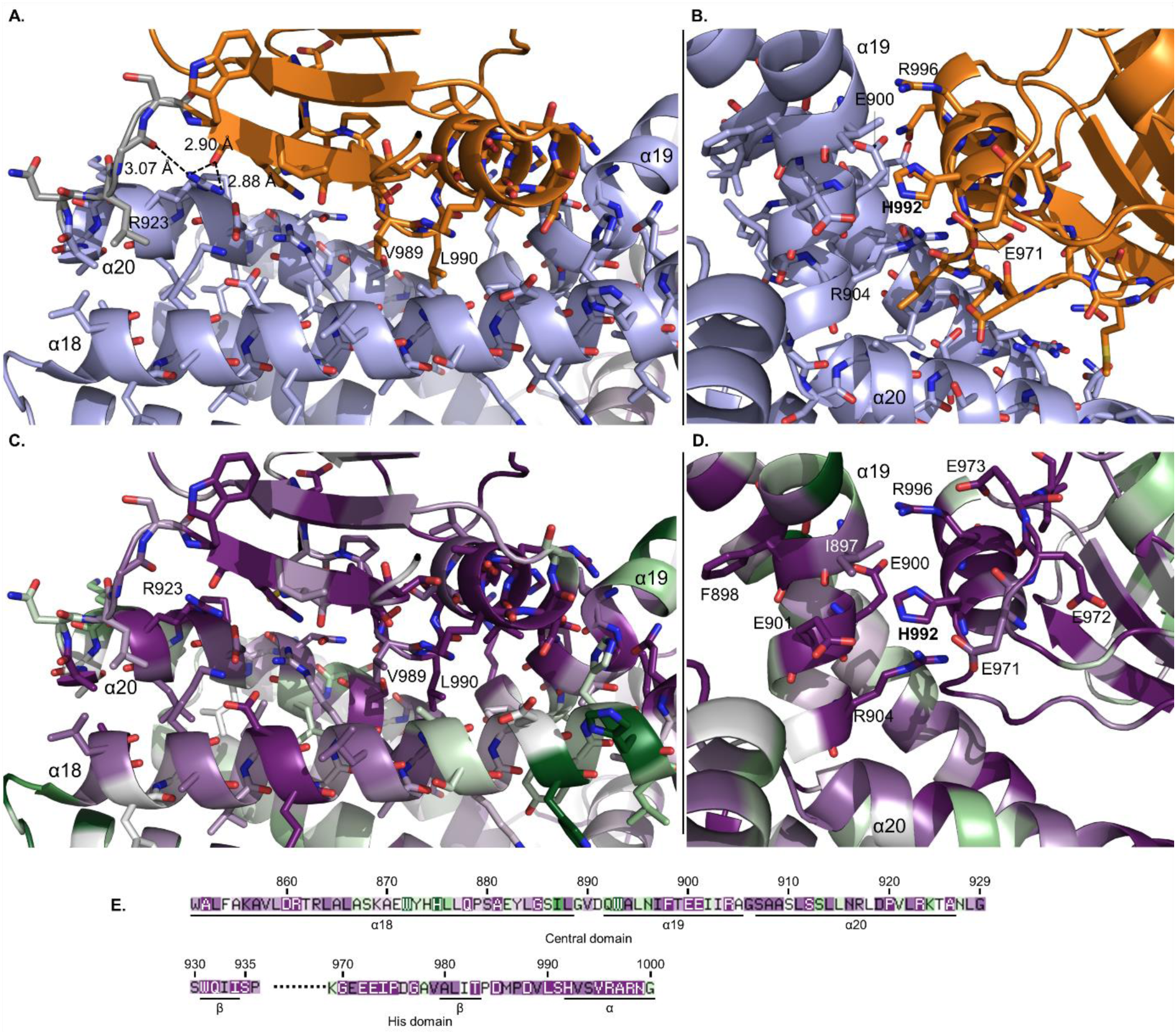
Interface between the structure of the central domain and the modelled histidine domain of StGWD1 in the open form. (A) and (B) Contact surface between the histidine domain (orange) and the central domain (mauve). Linker region is shown in gray. (C), (D) and (E) Conservation levels of the amino acids at the histidine domain-central domain interface, generated using the Consurf server. Deep purple: conserved. White: average. Green: variable. (D) Conserved amino acids in the vicinity of H992 are shown as sticks.

Focusing on the region around the catalytic H992 (Figure 8B and 8D), conservation analysis shows the presence of charged amino acids (E900, E901, R904) that might be directly or indirectly involved in the phosphate transfer reaction to the glucan. In LmRPH the authors proposed that a glutamic acid (E667) transfers a proton to the catalytic H825 upon phosphate transfer to the rifampin, while the Arginine R666 acts as a stabilizing base along with a yet unidentified residue to deprotonate the attacking hydroxyl of the rifampin. More recently, the phenolic phosphate synthetase BsPPS from *Bacillus subtilis* was characterized as a dikinase phosphorylating a wide range of polyphenols (Hsu et al., 2025). Mutagenesis revealed the crucial role of D627, which is believed to act as the deprotonating residue, as well as H629 and H630, thought to stabilize the phosphate. Despite the very low homology between the central domain of StGWD1 and the substrate binding domains of LmRPH and BsPPS (which in contrast, are homologs), we searched for equivalent residues in StGWD1. In our model, no other histidine is located close enough to H992, but we can suggest that E900 and R904, both located in the helix facing the H992 on the central domain, could be the proton donor and the stabilizer base. Alternatively, E971 which probably also assist the H992 in the autophosphorylation reaction (Figure 7C) could also be the proton donor, again in pair with R904 as stabilizer base.

## 4. Conclusion

By integrating biochemical characterization, AF2 modelling, X-ray crystallography and SAXS-driven modelling, we investigated the structure–function relationships of StGWD1, revealing a flexible protein composed of five distinct domains. At the N-terminus, StGWD1 carries two CBM45 domains, known to interact with starch, separated by a long partially folded linker. We solved the structure of the central domain by X-ray crystallography in two distinct conformations. The subsequent histidine and ATP-binding domains share conserved folds and residues with their counterparts in other dikinases. SAXS analysis proposed a model for the solution structure of the enzyme and strongly suggest a swiveling motion of the His domain between the ATP-binding and central domains. Regarding the role of the CBMs, our findings support a mechanistic model in which CBM45_1 would facilitate substrate targeting and approach, with the help of its flexible linker. For catalysis, the enzyme core must be positioned close to the glucan O6 hydroxyl groups and CBM45_2, located nearer to the central domain, would stabilize the latter in a precise orientation for phosphate transfer. In this configuration, glucan chains would access the central domain to react with the phosphorylated catalytic histidine, assisted by yet unidentified residues. However, the precise position and mode of glucan binding still remain unknown, especially given the two distinct conformations of the central domain.

Overall, our data provide a comprehensive view of StGWD1 architecture, highlight the dynamic features of its catalytic mechanism, and allow the identification of candidate residues potentially critical for activity. Future structural studies of larger StGWD1 constructs in complex with maltooligosaccharides, using X-ray crystallography or cryo-EM, combined with mutagenesis and biochemical analyses, are forthcoming to resolve these interactions in detail. This will be of primary interest for the understanding of StGWD1 mechanism and to drive enzyme engineering efforts aimed at enhancing phosphorylation levels or increasing the diversity of α-glucan substrates targeted by this intriguing enzyme.

## 5. Materials and Methods

### Full-length StGWD1 sequence and plasmids

Sequence of StGWD1 (Genbank Accession number Q9AWA5) without its signal peptide (aa 1-77) was codon-optimized for an expression in *E. coli* and ordered from TWIST bioscience (San Francisco, USA). Numbering of the protein started at the valine of the MVLTT N-terminal sequence. DNA fragment was inserted in a pET53 destination vector, using the Gateway technology to obtain pET53 His_StGWD1_Strep. The N-terminal 6His-tag was subsequently removed using by PCR (primer for: AGGAGATATACCATGGTCCTGACCACCGATACTAG and rev: ATCGGTGGTCAGGACCATGGTATATCTCCTTCTTAAAG to obtain pet53_STGWD1_Strep. A second PCR enabled the introduction of a Stop Codon to remove the C-terminal Strep-tag and obtain pet53_StGWD1 plasmid used for the enzyme production (primer for: CCCGTCCTCAAATGTAAGGTGGGCGCG and rev: GGTCGGCGCGCCCACCTTACATTTGAGGAC.

### Truncated forms and plasmids

DNA fragments corresponding to the desired truncated forms were generated by PCR and inserted in a pET28 vector. T7-peptide located at N-terminal end, and 6His-tag located at C-terminal end were removed, leading to proteins only fused to a 6His-tag at N-terminal end (with thrombin cutting site). First amino acid of the truncated form (indicated on Figure 3) was fused directly after the last one of the thrombin cutting site.

### Production of enzymes

*E. coli* BL21 (DE3) star cells were transformed with the desired plasmid. Precultures were grown overnight at 37 °C in liquid LB medium supplemented with 100 µg.ml^-1^ ampicillin or kanamycin depending on the plasmid used. Cultures in classic ZYM medium (Studier, 2005) supplemented in appropriate antibiotic were inoculated at OD_600nm_ = 0.05, and grown for 24 hours at 23 °C under 140 rpm agitation. Cells were harvested by centrifugation 15 min at 7 800 g, 6 °C. Cell pellets were stored at −20°C for one week maximum. Cells were resuspended in Binding Buffer (BB1 or BB2, see composition below) complemented with cOmplete™, Mini, EDTA-free Protease Inhibitor Cocktail, at OD_600nm_ = 100 and lysed by sonication on ice. The cell debris were separated from the clarified lysate (soluble fraction) by centrifugation for 25 min at 48 000 g at 6 °C.

### Purification of untagged StGWD1

Purification was carried out on an ÄKTA Xpress system (GE Healthcare), with a working flow of 3 ml.min^-1^. StGWD1 clarified and filtered lysate was loaded on a HiTrap Q HP 5ml anion exchange column equilibrated with BB1 (50 mM Tris/HCl, pH 7.5, 3 mM EDTA, 2.5 mM DTT and 10 % (w/v) glycerol). Column was washed with 7 CV of BB1, and StGWD1 was eluted with a two steps gradient of elution buffer 1 (EB1, BB1 + 1 M KCl): 0 to 4 % EB1 in 8.5 CV followed by 4 to 30 % EB1 in 21 CV. Fractions containing StGWD1 were collected, concentrated on Amicon centrifugal filter units (Millipore, 50 kDa cut-off) and injected into a size exclusion HiLoad 16/600 Superdex 200 pg column (GE Healthcare), equilibrated with 50 mM Tris/HCl, pH 7.0, 150 mM KCl and 2.5 mM DTT. Fractions containing full-length StGWD1 were collected, concentrated and freshly used. Calibration standards (ferritin, aldolase, conalbumin and carbonic anhydrase, from Cytiva) were injected in the same conditions.

### Purification of truncated forms

Purifications were carried out on an ÄKTA Xpress system (GE Healthcare), with a working flow of 1 ml.min^- 1^. Clarified lysate of truncated forms were loaded on a HisTrap HP 1 ml column equilibrated with BB2 (50 mM HEPES/KOH, pH 7.5, 6 mM MgCl_2_, 500 mM KCl, 20 mM Imidazole and 10 % (w/v) glycerol). Column was washed twice with 15 CV of BB2 and 5 CV of 97.5 % BB2/2.5 % EB2 (same as BB2 with 500 mM Imidazole). Proteins were eluted with a gradient of EB2 from 2.5 % to 50 % in 20 CV. Fractions containing the proteins of interest were concentrated and gel filtrated as the full-length enzyme in the same buffer.

### SDS-PAGE analysis

Samples (final dilution 20 times) were mix with loading buffer (commercial Laemmli Sample buffer (Biorad) supplemented with 2-Mercaptoethanol), denateured and loaded onto NuPAGE 3-8 % Tris-acetate precast gels. Migration was performed in commercial Tris/acetate/SDS buffer (Life technologies) for 1 hour at 150 V. Proteins were visualized by staining with PageBlue Protein Staining Solution (Thermo fisher) and visualization on a Gel Doc EZ System (Biorad).

### Activity assay

Crystalline maltodextrins were prepared as described by Hejazi et al., using maltodextrins dextrose equivalent 4-7 (Sigma) as substrate (Figure S17) (Hejazi et al., 2008). After SEC, StGWD1 fractions were prepared at 2.02 g.l^-1^ by dilution or concentration on Amicon (Millipore, 50 kDa cut-off). Protein concentration was estimated by absorbance at 280 nm using a NanoDrop instrument. StGWD1 activity was assayed in 50 mM HEPES/KOH pH 7.5, in presence of 6 mM MgCl_2_, 0.4 g.l^-1^ BSA and 0.5 mM DTT. Standard conditions were 1 mM ATP, 1 g.l^-1^ crystalline maltodextrins and 0.1 g.l^-1^ StGWD1. Reactions were carried out at 30 °C, in 2 ml tubes under 700 rpm agitation (in thermomixer, tubes also contained small magnetic stirrer to prevent sedimentation of insoluble material). Reactions were started by addition of the 20 X enzyme solution. Aliquots were withdrawn periodically and incubated 10 min at 55 °C to stop the reaction with minimal ATP degradation.

Truncated forms were assayed in the same conditions except that reaction buffer was 25 mM HEPES/KOH pH 7.5, in presence of 10 mM MgCl_2_, 0.4 g.l^-1^ BSA and 0.5 mM DTT. A protein molar concentration equivalent to that of StGWD1 was used (670 nM), and StGWD1 was also assayed in these conditions as a control.

### Determination of ATP, ADP and AMP concentrations by HPLC

After heating, samples were centrifuged 5 min at 13 000 g, and were analyzed without further dilution. 10 µl were injected on a C18 Zorbax-SB (250*4.6 mm, 5 µm, Agilent) column equipped with a C18 guard column (SecurityGuard Cartridges, 4*3 mm, Phenomenex) maintained at 24 °C and equilibrated with 80 mM KH_2_PO_4_/KOH pH 6.0 and 1 mM EDTA. Compounds were eluted with acetonitrile (ACN), with the following linear gradient: 0 min: 0 % ACN, 2 min: 0 % ACN, 9 min: 17.5 % ACN, 9.01 min: 0 % ACN, 13 min: 0 % ACN. Working flow rate was 1 ml.min^-1^, and it was raised to 1.3 ml.min^-1^ from 9 to 13 min. Compounds were quantified at 260 nm using a UV detector (Thermo ultimate 3000) and standard concentrations of 10, 50, 100, 500 and 1000 µM.

### Maltodextrins-P detection by mass spectrometry

Prior to analysis, samples were heated 5 min at 95 °C to precipitate the proteins, centrifuged 5 min at 13 000 g and analyzed without dilution. The analyses were carried out on an IC-MS platform of a capillary liquid anion exchange chromatography Thermo Scientific Dionex™ ICS-5000+ Reagent-Free™ HPIC™ system, coupled to a Thermo Scientific™ Q Exactive™ Plus hybrid quadrupole-Orbitrap mass spectrometer equipped with an electrospray ionization probe (ESI Ion Max).

Capillary liquid anion exchange chromatography was performed with the Thermo Scientific Dionex™ ICS-5000+ Reagent-Free HPIC system equipped with an eluent generator system (ICS-5000+EG, Dionex) for automatic base generation. Analytes were separated within 50 min, using a linear KOH gradient elution applied to a Thermo Scientific™ Dionex™ IonSwift™ MAX-100 column (250 x 0.25 mm) equipped with a guard column (50 x 0.4 mm, Dionex) at a flow rate of 0.12 µl.min^-1^. The gradient program was following: 0 min: 1 mM KOH, 0.01 min: 1 mM, 11 min: 5 mM, 20 min: 25 mM, 35 min: 100 mM, 40 min: 100 mM, 40.1 min: 1 mM, 50 min: 1 mM. The column and autosampler were thermostated at 35°C and 4°C, respectively. The injected sample volume was 1 µl.

Mass detection was carried out in a negative electrospray ionization (ESI) mode at a resolution of 70 000 (at 400 m/z) in full-scan mode, with the following source parameters: the capillary temperature was 350 °C, the source heater temperature, 350 °C, the sheath gas flow rate, 35 a.u. (arbitrary unit), the auxiliary gas flow rate, 9 a.u., the S-Lens RF level, 65 %, and the source voltage, 2.4 kV. A solution of IPA + 0.1 % (v/v) acetic acid is directly added in the source at a flow rate of 10 µl.min^-1^. Data acquisition was performed using Thermo Scientific Xcalibur software. Phosphorylated maltodextrins were investigated by extracting the exact mass with a tolerance of 5 ppm.

### Determination of phosphate incorporation

Prior to analysis, samples were heated 5 min at 95 °C to solubilize the glucan material, and centrifuged 5 min at 13 000 g. Glucans were hydrolyzed in 2 M trifluoroacetic acid (TFA) at 95 °C for 2 h. Samples were dried and wash 3 times with repeated cycles of solubilization in water followed by drying. Samples were resolubilized in their initial volume, diluted 50 times and mixed with 1 volume of internal standard solution to perform isotope-dilution mass spectrometry (Wu et al., 2005). The standard used was obtained from uniformly ^13^C-labelled extract from *E. coli,* and used to perform the absolute quantification of glucose-6-phosphate.

The samples were analyzed on an IC-MS platform composed of a liquid anion exchange chromatography Thermo Scientific Dionex™ ICS-5000+ Reagent-Free™ HPIC™ system, coupled to a Thermo Scientific™ Q Exactive™ Plus hybrid quadrupole-Orbitrap mass spectrometer equipped with a heated electrospray ionization probe (HESI-II).

Liquid anion exchange chromatography was performed with the Thermo Scientific Dionex™ ICS-5000+ Reagent-Free HPIC system equipped with an eluent generator system (ICS-5000+EG, Dionex) for automatic base generation. Analytes were separated within 50 min, using a linear KOH gradient elution applied to an IonPac AS11-HC column (250 x 2 mm, Dionex) equipped with an AG11-HC guard column (50 x 2 mm, Dionex) at a flow rate of 0.38 ml.min^-1^. The gradient program was the following: 0 min: 7 mM, 1 min: 7 mM, 9.5 min: 15 mM, 20 min: 15 mM, 30 min: 45 mM, 33 min: 70 mM, 33.1 min: 100 mM, 42 min: 100 mM, 42.5 min: 7 mM and 50 min: 7 mM. The column and autosampler were thermostated at 25 °C and 4 °C, respectively. The injected sample volume was 15 µl.

Mass detection was carried out in a negative electrospray ionization (ESI) mode at a resolution of 70 000 (at 400 m/z) in full-scan mode, with the following source parameters: the capillary temperature was 380 °C, the source heater temperature, 380 °C, the sheath gas flow rate, 50 a.u. (arbitrary unit), the auxiliary gas flow rate, 5 a.u., the S-Lens RF level, 60 %, and the source voltage, 2.75 kV. Data acquisition was performed using Thermo Scientific Xcalibur software. Glucose-6-phosphate was determined by extracting the exact mass with a tolerance of 5 ppm.

### Determination of glucose concentration

After acid hydrolysis, washing and solubilization (see previous paragraph), samples were diluted 100 times. High Pressure Anion Exchange Chromatography (HPAEC) was performed on a Thermo Scientific Dionex™ ICS-6000 HPIC™ system, equipped with Pulsed Amperometric Detector (PAD). Samples were injected on a CarboPac PA100 (250 x 2 mm) column equipped with a CarboPac PA100 (50 x 2 mm) guard column, at a flow rate of 0.25 ml.min^-1^. Analytes were separated using a linear gradient of sodium acetate, in 150 mM NaOH. Gradient was programmed as following: 0 min: 5 mM, 30 min: 500 mM, 40 min: 500 mM, 40.01 min: 5 mM, 50 min: 5 mM. The column and autosampler were thermostated at 30 °C and 10 °C, respectively. The injected sample volume was 10 µl. Standard concentrations were 5, 10, 15, and 20 mg.l^-1^.

### Differential scanning fluorimetry

Differential Scanning Fluorimetry (DSF) was used to test the stabilization effect of carbohydrate ligands, as well ATP, ADP, AMP and AMP-PNP (Table S1) with final concentrations of 5 µM of enzyme and 10 X of SYPRO orange (supplied 5000 X, Sigma). A ramp from 20 to 80 °C was applied with a gradient ramp of 0.3 °C.s^-1^ on a C1000 Touch Thermal Cycler (Biorad). NanoDSF experiments were performed using a Tycho-NT6 apparatus (Nanotemper) with an enzyme concentration of 1 g.l^-1^ and various concentrations of Adenosine-5’-[(β,γ)-imido]triphosphate (AMP-PNP, Roche).

### Generation of AlphaFold2 models

AlphaFold2 (AF2) models were generated using the Google colab AlphaFold2_advanced, with the following parameters: all parameters by default, except use_turbo: false and max_msa: 256:512. To model ΔCBM2 with the His domain in the transfer conformation, we first generated an AF2 model for the construct ΔCBM2ΔATP. Then, we manually merged this model with the ATP-binding domain. The linkers between the central domain, the His domain and the ATP-binding domain have been re-modelled using the Robetta server (Song et al., 2013).

### Chymotrypsin treatment

StGWD1 was incubated with chymotrypsin at a molar ratio chymotrypsin:StGWD1 of 1:500, in StGWD1 SEC buffer supplemented with 1 mM CaCl_2_. Reactions were incubated at 4 °C without agitation. 4 µl aliquots were withdrawn periodically and the proteolysis reaction was quenched by addition of 1 µl of 25 mM phenylmethylsulfonyl fluoride (PMSF) before SDS-PAGE analysis.

### Transfer onto PVDF membrane for N-terminal sequencing

Aliquots from the digestion at 22 g.l^-1^ were diluted 4 times, and mixed with loading buffer (commercial Laemmli Sample buffer (Biorad) supplemented with 2-Mercaptoethanol, prepared 4 X) at a volume ratio 4:1. 15 µl were loaded on a SDS-PAGE gel, that was run as before. Transfer onto PVDF membranes (0.2 µm, Thermo Fisher) was performed on ice, in 10 mM CAPS pH 11 and 10 % (v/v) methanol, at 0.8 mA.cm-2 for 1h15. Bands were visualized with Ponceau Red, cut out of the membrane, and then subjected to N-Terminal sequencing by automated Edman degradation, using a Shimadzu PPSQ 31B sequencer.

### Crystallization trials and data collection

Commercial screens from Qiagen (AmSO_4_, Classics, Compas, JCSG I, II, III and IV, JCSG+, Morpheus, Optisalts, PACT, PEG 1 and 2, pH Clear and ProComplex) as well as manually designed conditions were used to screen for crystallization conditions. Sitting drops were made by mixing 0.2 µl of enzyme solution with 0.2 µl of reservoir solution using a Mosquito robot (TTP Labtech). Several conditions led to crystals of the central domain, and the best crystal for the open form was obtained in 2 M NaCl, 10 % PEG 6K with a protein concentration of 10 mg.ml^-1^ after 2 weeks of incubation at 12 °C. The crystal for the closed form was obtained in 0.7 M tri-Na citrate, 0.1 M HEPES-Na, pH 7.5. All crystals were harvested and cryo-protected in 20 % ethylene glycol mixed with the crystallization solution before cryo-cooling into liquid nitrogen. Diffraction data were collected at beamline ID23-1 of the European Synchrotron Radiation Facility (ESRF), Grenoble.

### Structure solution, refinement and validation

Data sets were integrated using XDS and further processed with the programs contained in the CCP4 suite (Agirre et al., 2023). Structure for the central domain were solved by molecular replacement using Phaser (Read, 2001) and the AlphaFold2 model of the central domain as search model. Structures were automatically rebuilt using Buccaneer (Cowtan, 2012) and submitted to restrained refinement using Refmac (Murshudov et al., 1997) alternated with manual rebuilding cycles using Coot (Emsley et al., 2010). Data collection and refinement statistics are show in Table S2. Final models were validated using the Molprobity (Williams et al., 2018) and deposited in the PDB with accession number 9HA9 and 9QVI.

### Small Angle X-ray Scattering (SAXS)

SAXS data were collected at beamline BM29 of the European Synchrotron Radiation Facility (Grenoble, France). Freshly purified proteins were concentrated to 10-20 mg/mL using an Amicon centrifugal filter (Millipore, 10 – 50 kDa cut-off). Samples (50 µL) were injected into a size exclusion column (S200 Increase 3.2/300, Cytiva) and eluted inside the SAXS cell (set at 20°C) using gel filtration buffer (50 mM Tris/HCl, pH 7.0, 150 mM KCl). A photon energy of 12.5 keV (0.99 Å) was used with a beam size at the sample location of approx. 200 x 200 µm. The Pilatus3 x 2M detector was located at 2.74 m from the sample, giving an accessible Q-range from 0.006 to 0.55 Å^-1^. 2D-data integration and reduction to 1D scattering profiles was automatically performed by the BM29 data processing software (Tully et al., 2023). Further processing was done using the ATSAS suite (Manalastas-Cantos et al., 2021). In particular, CHROMIXS was used to select, subtract and average the HPLC frames. Radii of gyration were determined by Guinier approximation using PRIMUS and Dmax was calculated using the GNOM software. Ab-initio molecular envelops were calculated with GASBOR and rigid-body molecular modelling was performed using the DADIMODO webserver (Rudenko et al., 2019). In all the models reconstructed, the central domain and the ATP-binding domain were considered as a single object (not able to move one from another). Fitting (χ²) of the DADIMODO models against the experimental data was recalculated with CRYSOL.

The SAXS measurements for the AMP-PNP binding experiment were performed at Laboratoire de Génie Chimique, Toulouse, on a XEUSS 2.0 instrument (Xenocs, Grenoble, France) equipped with a Genix3D source that produces a X-ray beam (8 keV and 30.10^6^ ph.s^−1^) providing a size resolution of approximately 0.7 × 0.7 mm. Freshly purified ΔCBM45_2 was concentrated to 10 mg/mL using an Amicon (Millipore, 10 - 50 kDa cut-off) and analyzed in the presence/absence of 2 mM AMP-PNP + 6 mM MgCl_2_. Sample aliquots (50 µL) were transferred from the sample holder (maintained at 10 °C) to the measurement cell placed under vacuum to limit air absorption. Data were collected on a 160 × 160 mm area DECTRIS detector (Pilatus 1M) at a sample-detector distance of 1.216 m, thus procuring a measurement range from 0.005 to 0.5 Å^-1^. Each sample dataset is an average of 6 measurements with a data collection time of 1800 s. Data integration and reduction were performed using the FOXTROT software. Data were then averaged and buffer subtracted as described above. The OLIGOMER software (Konarev et al., 2003) was used to determine the contribution of the transfer and the ATP-cleavage conformations to each set of SAXS data.

## Supplementary materials

Supplementary material file: Figures S1 to S16 and Tables S1 to S3

## Author contribution

Conceptualization MRS, CM, GC, TL

Formal analysis TL, NC, GC, DG, CM, MRS

Investigation TL, NC, GC

Visualization TL, GC

Methodology TL, NC, DG, ES, PM, PR, GC, CM, MRS

Funding acquisition CM,

Supervision CM, GC, MRS

Writing – original draft TL

Writing – review & editing: TL, GC, PR, CM, MRS

## Acknowledgments

The authors thank the Integrated Screening Platform of Toulouse (PICT, IBiSA) for accessing protein purification, analytical and crystallization equipment. They are grateful to Sandra Pizzut and Sophie Bozonnet for providing technical assistance on the PICT-ICEO facility of TBI, which is part of PICT and a member of IBISBA-FR (https://doi.org/10.15454/08BX-VJ91), the French node of the European research infrastructure, IBISBA (www.ibisba.eu). They also thank the MetalToul platform of TBI for giving access to the LC-HRMS and LC-HRMS/MS facility, the Federation FERMaT, Université de Toulouse, France for accessing the in-house SAXS instrument, and the ESRF (Grenoble, France) and ALBA (Barcelone, Spain) synchrotrons for accessing crystallography and SAXS beamlines.

## Data availability statement

Data available on request from the authors.

## Abbreviations

AF2: AlphaFold2
AMP-PNP: Adenylyl-imidodiphosphate
CBM: Carbohydrate Binding Module
G6P: Glucose-6-phosphate
GWD1: Glucan Water Dikinase 1
PPDK: Pyruvate, Phosphate Dikinase
PEP: Phosphoenolpyruvate

